# Faster R-CNN Model for Predicting Endometrial Cancer Recurrence: A Five-Subclass Approach with Texture Analysis and Validation

**DOI:** 10.1101/2025.06.13.659644

**Authors:** Huy Duc Vu, Hui Joong Lee, Juhun Lee, Hyun Jung Lee, Sung Won Youn

## Abstract

**Purpose:** This study investigates the prediction of endometrial cancer (EC) recurrence using MRI-based texture analysis and deep learning. EC recurrence remains a major clinical challenge, and accurate prognostic tools are needed. The study aims to identify significant imaging biomarkers for recurrence risk and evaluate the performance of a Faster R-CNN deep learning model in classifying recurrence-free survival (RFS) and recurrence (REC) groups.

**Materials and Methods:** A total of 112 patients with histopathologically confirmed EC who underwent pre-treatment MRI between 2009 and 2018 were included. MRI texture analysis was performed using histogram and gray-level co-occurrence matrix (GLCM) methods, analyzing perimeter, integrated density, entropy, and homogeneity. A Faster R-CNN deep learning model was trained and tested on five image subclasses, using 20,000 iterations to optimize performance. REC and RFS were classified based on clinical outcomes, and probability scores were calculated to define the REC score. Feature importance was assessed using a Random Forest model.

**Results:** Significant clinical predictors of recurrence included high-grade tumors, FIGO stage III-IV, and cervical invasion. Texture analysis revealed notable differences in perimeter, integrated density, contrast, entropy, and homogeneity between REC and RFS groups. Faster R-CNN achieved an average precision of 0.86 and an AUC of 0.61. Feature importance analysis identified area as the most significant factor, while REC_score was independent of texture-based parameters.

**Conclusion:** MRI texture analysis combined with deep learning provides valuable predictive insights for EC recurrence. The study highlights the potential of integrating machine learning with imaging biomarkers for personalized risk assessment, offering a promising approach for improving EC patient management..

## Introduction

Endometrial cancer (EC) is one of the most prevalent gynecological malignancies worldwide, with an increasing incidence over the past decades[1]. Although early-stage EC generally has a favorable prognosis, recurrence remains a significant concern, particularly in patients with high-grade tumors and advanced-stage disease. Accurate patient identification at high risk for recurrence is essential for improving treatment strategies and optimizing clinical outcomes. Traditional prognostic factors, including histopathological features, FIGO staging, and molecular markers, have been widely used for recurrence risk assessment [2]. However, these factors alone may not sufficiently capture tumor heterogeneity and variations in treatment response, necessitating the development of more advanced predictive models [3]. Among preoperative diagnostic tools, magnetic resonance imaging (MRI) plays a crucial role in EC evaluation, providing detailed anatomical and functional insights into tumor characteristics, depth of myometrial invasion, cervical stromal involvement, and lymph node metastasis [4, 5]. Experienced radiologists analyze MRI images to predict the stage and prognosis of EC by examining various features like the tumor’s size, location, and extent of invasion [5].

Recent advancements in deep learning technology have led to remarkable improvements in medical imaging analysis [6]. In particular, Faster R-CNN has been widely adopted across various medical fields due to its superior object detection performance [7]. By integrating Region Proposal Networks (RPN), Faster R-CNN enhances both efficiency and accuracy compared to previous models, making it highly effective for lesion detection and characterization in oncologic imaging. Deep learning-based convolutional neural networks (CNNs) have the potential to identify complex imaging patterns associated with EC, offering novel capabilities in lesion detection and prognosis evaluation [8]. Furthermore, comparing deep learning models with traditional radiomic approaches, such as texture and histogram analysis, provides deeper insight into the advantages and limitations of artificial intelligence (AI) in medical imaging. Unlike conventional radiomic techniques that rely on predefined feature extraction, Faster R-CNN automatically learns intricate imaging patterns linked to EC prognosis, minimizing human bias and enhancing interpretability [9, 10].

This study aims to validate the Faster R-CNN model for MRI-based EC recurrence prediction. Specifically, we evaluate the performance of Faster R-CNN in analyzing MRI perfusion maps and compare its predictive accuracy with traditional texture-based methods. We hypothesize that the Faster R-CNN-derived quantitative objective index will provide independent prognostic value beyond conventional texture features. This numerical metric allows direct comparison with existing imaging analysis methods, offering a standardized framework for EC recurrence risk assessment. Additionally, we aim to assess the robustness of Faster R-CNN in detecting EC recurrence and explore its clinical applicability in oncologic imaging.

To ensure the effectiveness of deep learning-based classification, we rigorously validate the Faster R-CNN model in predicting EC recurrence risk. Its performance is compared against traditional texture and histogram analysis to establish a robust validation framework. This study aims to bridge the gap between experimental AI-driven models and their practical implementation in EC management.

Thus, the primary objective of this study is to validate the feasibility of a Faster R-CNN-based MRI deep learning algorithm for EC recurrence prediction, comparing it with conventional histogram and texture analysis of pelvic MRI. By assessing its predictive reliability and generalizability, we aim to establish a robust evaluation framework for deep learning-based medical imaging analysis. If validated, our approach may serve as a valuable tool for early recurrence detection, guiding treatment planning and personalized patient management in EC care.

## Materials and methods

### Institutional review board approval

The institutional review board of Kyungpook National University Hospital (KNUH 2017-06-012) approved the study. Due to the retrospective nature of the study, The institutional review board of Kyungpook National University Hospital waived the need of obtaining informed consent. All methods were performed in accordance with relevant guidelines and regulations.

### Study Population

The study included 112 patients diagnosed with EC who underwent pre-treatment MRI between June 2009 and March 2018. All patients fulfilled the following inclusion criteria: (1) histopathologically confirmed as endometrial cancer; (2) available multi-parametric gadolinium-enhanced pre-treatment MRI including sagittal, axial, and/or coronal plane before any treatment, and (3) complete surgical resection of the primary tumor with documented clinical follow-up. Patients with incomplete MRI scans or loss to follow-up were excluded.

### Clinical endpoint

Patients were followed up with clinical evaluation, cytological assessment and imaging studies every three months for the first two years. In the third, fourth, and fifth years, follow-ups occurred every three months for the first two years, every six months during the third to fifth years, and annually thereafter up to ten years. Recurrence (REC) was defined as presence of focal presence of intrapelvic tumor or lymphatic metastases of EC after treatment and confirmed by multiparameter contrast-enhanced imaging or histology. Recurrence-free survival (RFS) was defined as the time period between definitive treatment and first imaging evidence of recurrence. Abdominal computed tomography or ultrasound imaging was performed at regular intervals. Patients received images every three months for the first year after the procedure and every 6 months thereafter.

### MRI and perfusion mapping

All examinations were performed using a 3.0-tesla MRI machine (Skyra; Siemens Health Care, Erlangen, Germany). Prior to examination, patients underwent 6 hours of fasting followed by intramuscular administration of 20 mg of hyoscine butylbromide (Buscopan; Boehringer Ingelheim) to inhibit bowel movement. A set of 8-channel phased array body coils was applied, and MRI examination was performed using the same parameters for field of view, slice thickness, echo time, repetition time for T1-WI, and contrast-enhanced T1-WI acquisition. A normalized perfusion map was generated by first subtracting the non-contrast-enhanced image from the contrast-enhanced image, and then dividing the result by the contrast-enhanced image. This standardization ensured consistency in MRI intensity values across different acquisitions (Figure 1). Each region of interest (ROI) was defined using the outline of the EC tumor at the level with the largest area of the tumor.

**Figure 1.**
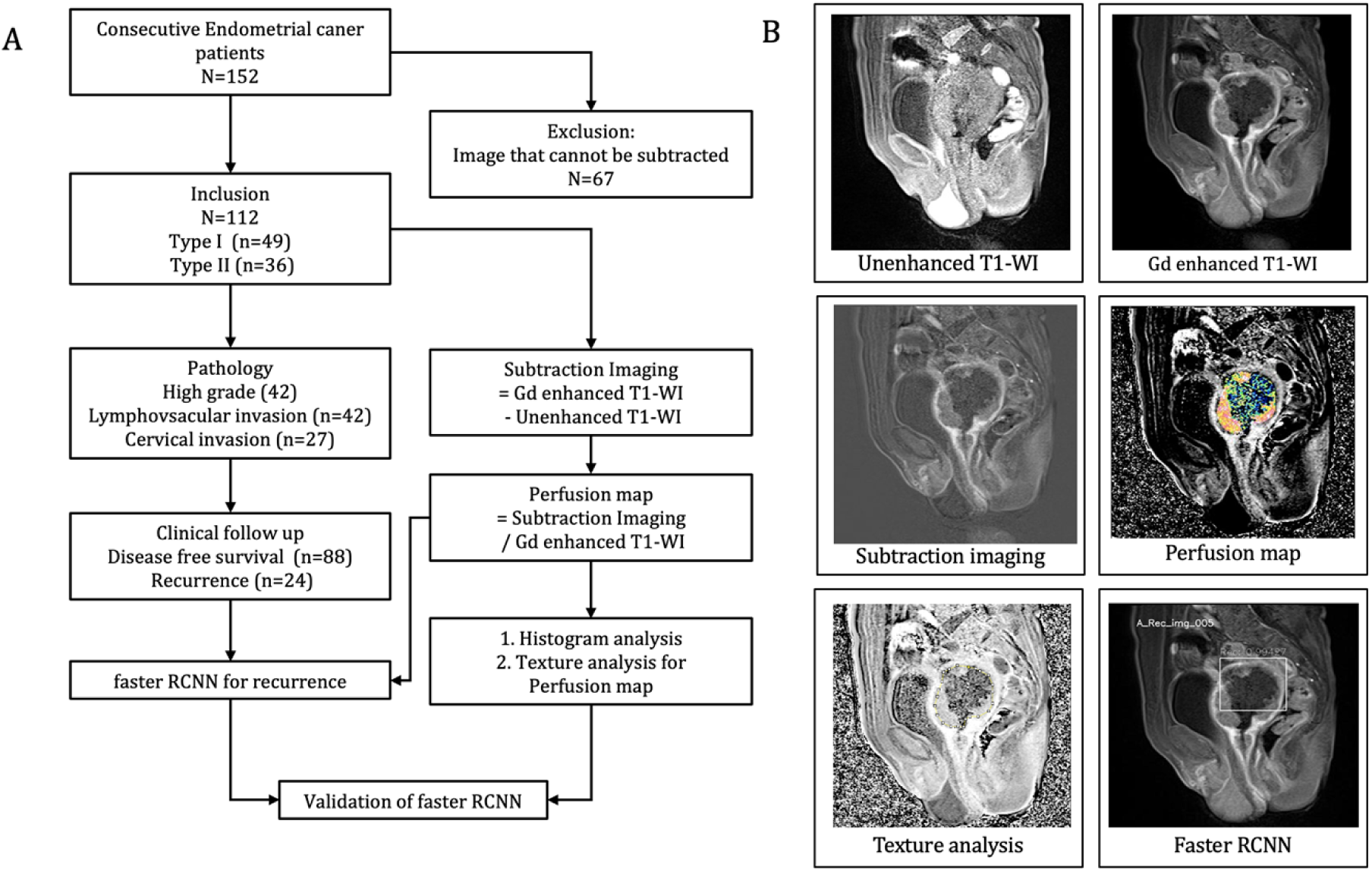
(A) Flow chart illustrating the selection process of study participants, including the inclusion and exclusion criteria applied to derive the final study cohort. (B) Image preparation process detailing the steps for preprocessing and analyzing images. A non-contrast-enhanced image was subtracted from the contrast-enhanced image to generate a subtraction image, which was then divided by the contrast-enhanced image to obtain a normalized perfusion image. The region of interest (ROI) for endometrial cancer was defined on the perfusion map. Image data were imported into a web-based VGG Image Annotator tool, where a minimal rectangular ROI containing the endometrial cancer was manually drawn.

### Texture analysis GLCM

Texture analysis was performed using a gray-level co-occurrence matrix (GLCM), the most common and sensitive texture descriptor to calculate lesion heterogeneity in greater detail from the texture data [13]. The area, mean and standard deviation of the signal intensity, and integrated density of the affected ovary with endometrioma were compared with those of the normal ovary. Integrated density, as the sum of all pixel intensities in the ROI, indicates the total amount of contrast enhancement effect in this study. The parameters included in the angular second moment (ASM), contrast, correlation, contrast, correlation, inverse difference moment (IDM), and entropy for the same ROI in the perfusion map, were calculated using the GLCM plugin using ImageJ (1.50i, National Institutes of Health, Bethesda, MD, USA, https://imagej.nih.gov/ij).

### Deep learning

#### 1. Computing environment

Deep learning was performed using a central processing unit (Intel® Core ™ i7-8700K, 12 cores 3.70 GHz), a graphics processing unit [Geforce RTX™ 2080 Ti 12 GB (NVIDIA®, Santa Clara, CA, USA)], an Ubuntu 18.04 operating system (Canonical Ltd., London, UK), CUDA 10.1 computing environment (NVIDIA®, Santa Clara, CA, USA), TensorFlow 1.12, and Python 3.6.

#### 2. Structure of Faster R-CNN

The structure of the faster R-CNN has been described in the previous literature [7]. Briefly, the faster R-CNN consisted of the RPN and the Fast R-CNN (Figure 2). Using the input images, the RPN extracted the feature map that was fed into the backbone convolutional neural network. The Faster R-CNN uses the ResNet-101 model to extract features from the perfusion map.

**Figure 2.**
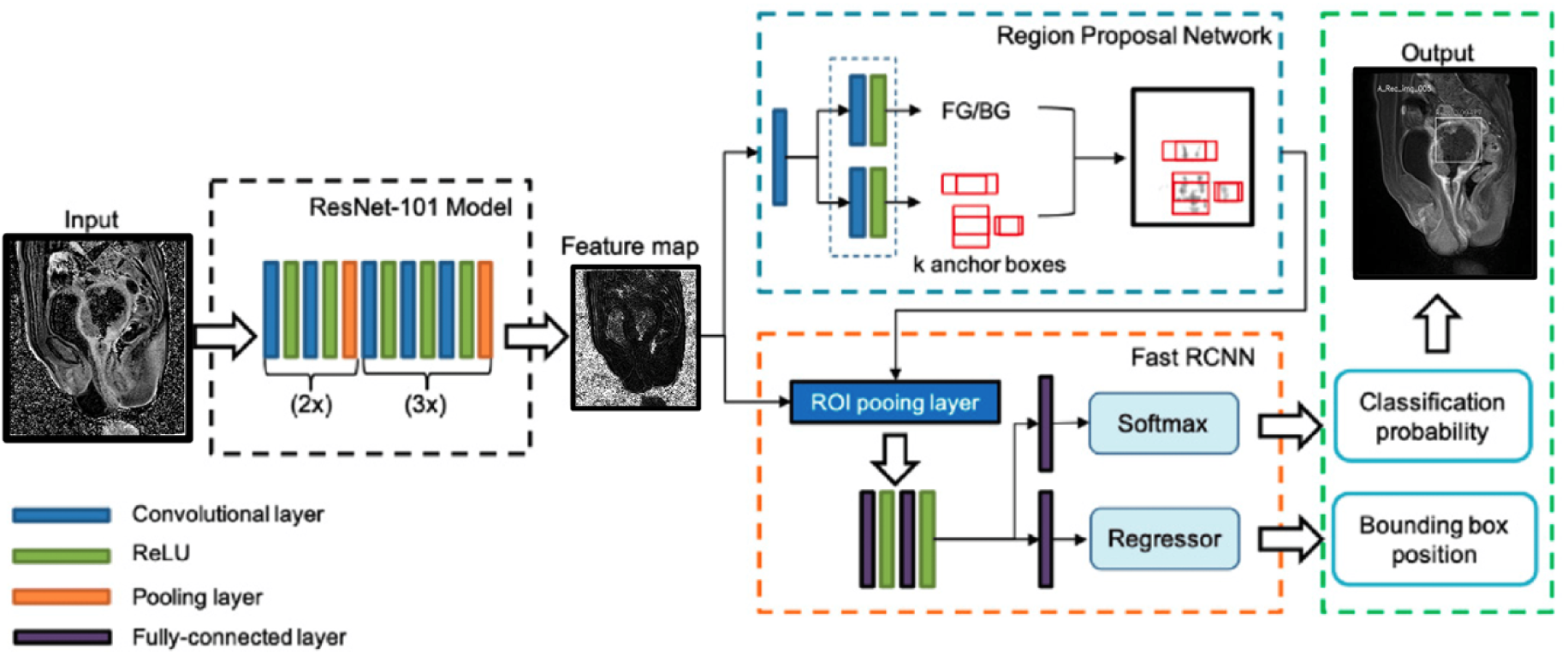
Architecture of faster region-based convolutional neural network (faster R-CNN). From the input image, a ResNet-101 model without fully connected layers extract a feature map. The RPN is responsible for determining the presence of objects in the input image by placing a series of anchors. These anchors are evaluated against each pixel in the feature map, determining their objectness score and refining their coordinates as rectangular regions of interest (ROI) by using rectified linear unit (ReLU) activation function. The output features from the ROI pooling layer are then passed through fully connected layers with softmax and regressor branches. This process ultimately produces the classification probability and bounding box position.

The RPN learns every point in the output feature map to determine whether an object is present on the input image at the corresponding location by placing a set of anchors on the input image for each location on the feature map. As the network propagates each pixel in the feature map, these anchors are checked to determine the objectness score to refine the anchor’s coordinates of the bounding boxes as the ROI. The Fast R-CNN detector also consists of a CNN backbone, an ROI pooling layer, and fully connected layers followed by two branches for classification probability and bounding box regression. The bounding box proposals from the RPN are used to pool features from the backbone feature map implemented by the ROI pooling layer. The ROI pooling layer works by taking the region corresponding to a proposal from the backbone feature map, dividing this region into a fixed number of sub-windows, and performing max pooling over these sub-windows. Finally, the output features from the ROI pooling layer are fed into the fully connected layers and the softmax and bounding box branches.

#### 3. Image classification and annotation

In this study, imaging data including perfusion map were classified into recurrence or RFS according to the patient’s clinical outcome. For every patient, one image containing most distinctive feature was extracted from the PACS and saved as a JPEG file. A board-certified radiologist imported image data into a web-based VGG Image Annotator tool [21] and manually drew a minimal rectangular ROI containing EC. After completing localization of the rectangular ROIs, a raw comma-separated values (CSV) file containing the bounding box coordinates (x and y, width, and height) was created. The bounding box coordinates in the raw CSV file were converted into the required format (x, y, x + width, and y + height) with the pattern label identified for each image, to be read by the Python-based Faster R-CNN pipeline.

#### 4. Dataset construction

After image classification and data annotation, the datasets were divided into five subclasses, as follows; subclass A (total; recurrence, RFS = 24;20,4), subclass B (22;17,5), subclass C (22;17,5), subclass D (22;17,5), and subclass E (22;17,5), respectively.

#### 5. Training and testing of faster R-CNN

To determine training iteration numbers, all the images in the dataset was trained preliminarily with iterations of 5000, 10,000, 20,000, and 50,000 steps to find optimal training epoch. These preliminary training results were then presented as mean average precision (mAP) values and the AUC-ROC. Variations in the mAP at the intersection over the union in the range of 50–95% and the change trend in the loss function values at varied iterations were measured and the condition showing the highest mAP was selected for main training and testing.

One subclass was considered as the test set, with the remaining serving as the training dataset. A 4:1 ratio cross validation method was used to train the DL model to predict recurrences (AL^REC^) using the images and annotations from four subclasses. These were then tested using the remaining subclass, and this process was repeated five times for each subclass.

In the test set, the Tensorflow-Phython library reported multiple probability value of each candidate region, and printed the highest probability value of candidate region with its predicted label such as probability of recurrence (Prob*^REC^*) and probability of RFS (Prob*^RFS^*) with range 0–1.

The highest probability value with predicted label was used to undergo ROC analysis. There were two labels: recurrence and RFS, and each label was inferred simultaneously by trained model at the input of same test set image. To differentiate recurrence feature from those of RFS, a new score system (REC score) was defined and calculated as follows:

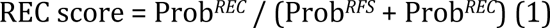

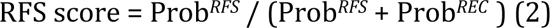

where Prob*^REC^*, and Prob*^RFS^* represent the recurrence and non-recurrence probabilities predicted by Faster R-CNN, respectively.

### Statistical analyses

Statistical analyses were performed using SPSS Statistics Version 23.0 for Windows (IBM Corp, Armonk, NY). Paired *t-*test was used to examine differences in numerical variables from those in the texture analysis. Statistical significance was set at a *p-value* of <0.05. To evaluate the predictive performance of MRI-derived texture features and deep learning-based REC_score, we implemented a Random Forest classifier using R (version 4.2.1, https://CRAN.R-project.org). The area under the receiver operating characteristic curve (AUC–ROC) was used to evaluate the performance of texture analysis. Receiver operating characteristic (ROC) curve analysis was also used to assess the diagnostic performance of the Faster R-CNN, and the area under the curve (AUC), sensitivity, and specificity were calculated. The correlation matrix plot was created using packages “ellipse” and “corrplot” in R.

## Results

### I. Clinical characteristics of patients

This study included 112 patients with pathologically confirmed EC after hysterectomy. The mean age was 58.40 ± 9.88 years, andrecurrence was observed in 24 (21.4%) patients. Histology (p-value = 0.022), FIGO stage > III and IV (p-value = 0.007), lymphovascular invasion (p-value = 0.005), and invasion into the cervical gland (p-value = 0.007) and stroma (p-value = 0.005) were seen to be significantly associated with risk of recurrence (Table 1).

**Table 1.** Patient demographics of endometrial cancer of patients with or without recurrence.

|  | Total<br>N=112 | No<br>recurrence<br>N=88 | Recurrence<br>N=24 | *p*-<br>value |
| --- | --- | --- | --- | --- |
| Age, years | 58.40 ±<br>9.88 | 56.38 ± 9.58 | 65.83 ±<br>7.09 | <0.001 |
| Histology and grade - no.<br>(%) |  |  |  |  |
| endometrioid, grade 1 | 27 (24.1) | 25 (28.4) | 2 (8.3) | 0.022 |
| endometrioid, grade 2 | 32 (28.6) | 26 (29.5) | 6 (25.0) |  |
| endometrioid, grade 3 | 13 (11.6) | 12 (13.6) | 1 (4.2) |  |
| mixed | 18 (16.1) | 12 (13.6) | 6 (25.0) |  |
| serous | 8 (7.1) | 3 (3.7) | 5 (7.1) |  |
| clear cell | 6 (5.4) | 4 (4.5) | 2 (8.3) |  |
| others | 8 (7.1) | 6 (6.8) | 2 (8.3) |  |
| Figo stage - no. (%)* |  |  |  |  |
| I-II | 85 (75.9) | 72 (81.8) | 13 (54.2) | 0.007 |
| III-IV | 27 (24.1) | 16 (18.2) | 11 (45.8) |  |
| Lymphovascular<br>invasion | 42 (37.5) | 27 (30.7) | 15 (62.5) | 0.005 |
| Endocervix |  |  |  |  |
| gland invasion | 28 (25.0) | 16 (18.2) | 11 (45.8) | 0.007 |
| stromal invasion | 27 (24.1) | 15 (17.0) | 11 (45.8) | 0.005 |
| BMI, kg/m2** | 24.83 ±<br>4.87 | 24.85 ± 5.33 | 24.72 ±<br>2.56 | 0.875 |

\*Stages were assigned according to the International Federation of Gynecology and Obstetrics (FIGO) 2009 classification; stages range from I to IV, with higher stages indicating more advanced spread of cancer.
\*\*Body-mass index (BMI) is the weight in kilograms divided by the square of the height in meters.

In the multivariate Cox regression model, predictors of tumor recurrence was found; tumor recurrence was related to the high-grade tumor (HR:3.911; 95% CI, 1.699 to 9.007; p=0.001), advanced (III or IV) FIGO stage (HR:2.956; 95% CI, 1.321-6.612; p=0.008), endocervical gland (HR:2.992; 95% CI, 1.318-6.792; p=0.009) and stromal invasion (HR:3.619; 95% CI, 1.611-8.129; p=0.002). No such associations with lymphovascular invasion lymphovascular invasion (HR:2.066; 95% CI, 0.869-4.916; p=0.101) and lymph node metastasis (HR:1.818; 95% CI, 0.772-4.282; p=0.171) were seen (Figure 3).

**Figure 3.**
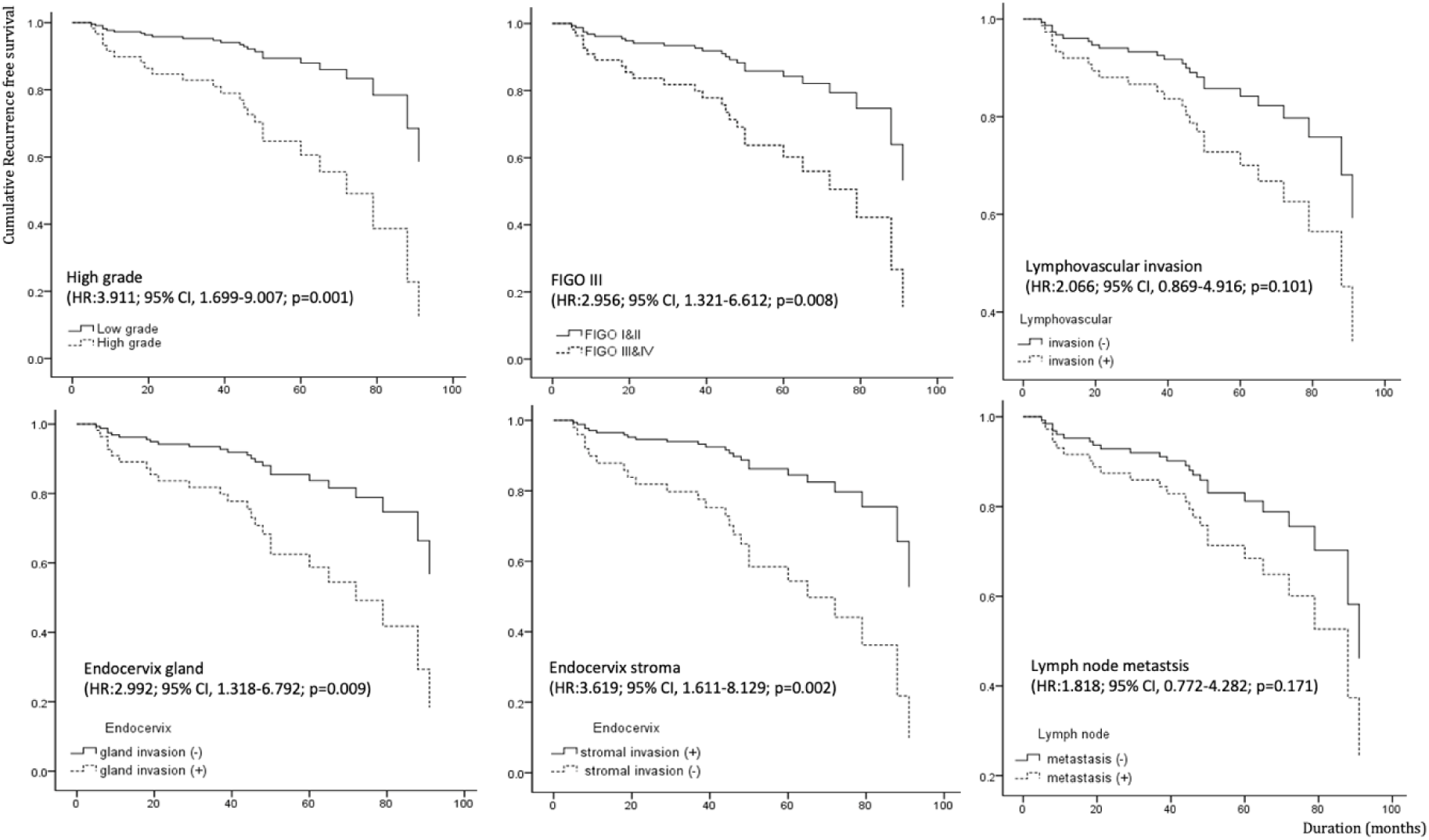
Cumulative recurrence-free survival (RFS) of endometrial cancer according to high-grade, FIGO staging, lymphovascular invasion, endocervical gland and stromal invasion and lymph node metastasis

**Figure 4.**
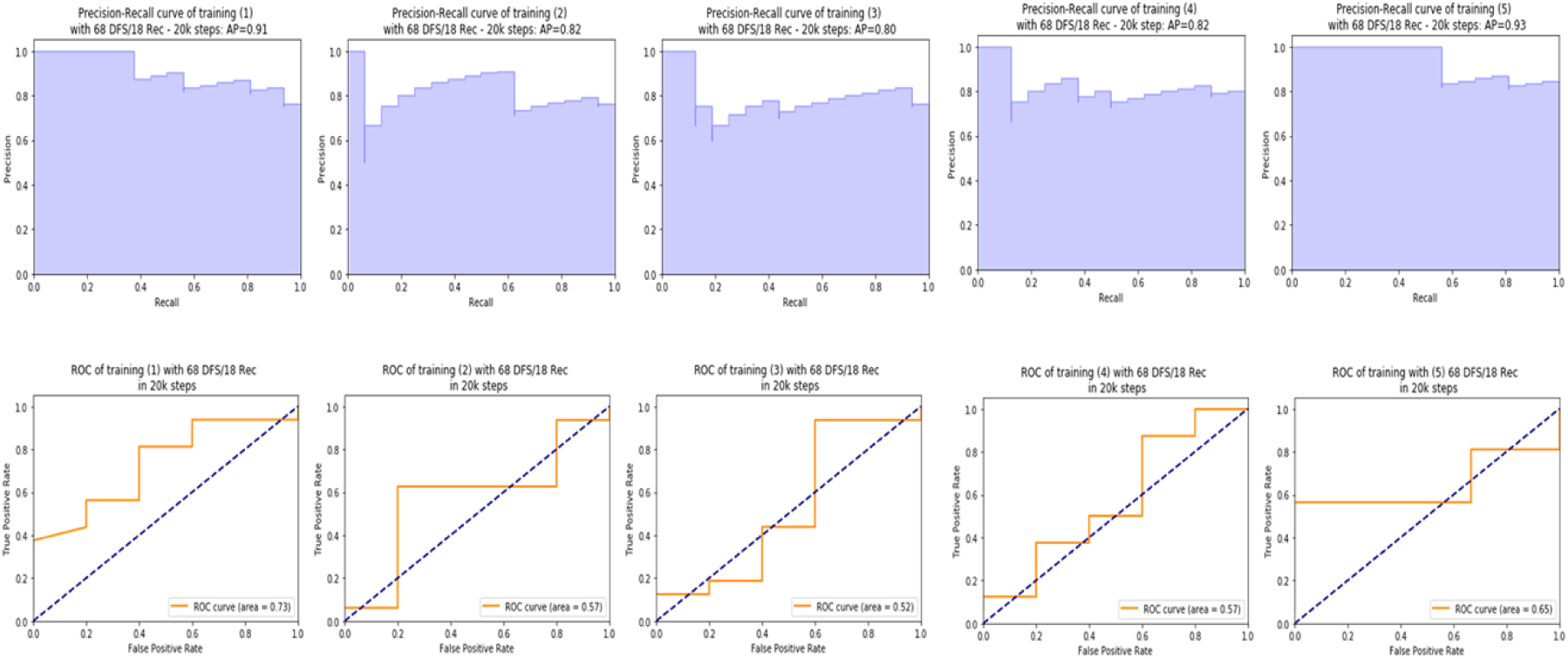
Precision-recall (top) and ROC (bottom) curves for five models trained on different subsets of the endometrial cancer dataset. A 4:1 training-to-testing ratio was used, with each training session including 68 Disease-Free Survival and 18 Recurrence cases, while testing was performed on 18 DFS and 5 Rec cases per subset. Precision-recall curves show model performance in recurrence prediction, with Average Precision values displayed. ROC curves illustrate the discriminative ability of the model, with area under the curve values indicating model discriminative power.

### II. Calculation of pattern score for each image using faster R-CNN

The 20,000 iteration was chosen for the training session and construction of the AL^REC+RFS^, at which highest mAPs and lowest loss function values were recorded (average precision: 0.93, AUC: 0.65). AUCs were obtained for each model’s testing results using the remaining four subclasses as training datasets. The average precision and AUC for subclass A (trained using subclasses B + C + D + E) were 0.91 and 0.73, respectively; for subclass B (trained using subclasses A + C + D + E) were 0.82 and 0.57, respectively; for subclass C (trained using subclasses A + B + D + E) were 0.8 and 0.52, respectively; for subclass D (trained using subclasses A + B + C + E) were 0.82 and 0.57, respectively; and for subclass E (trained using subclasses A + B + C + D) were 0.93 and 0.65, respectively. The overall mean precision and AUC (calculated by averaging the five test results) were 0.86 ± 0.06 and 0.61 ± 0.08, respectively (Figure4).

### III. Characteristics of imaging data

Histogram and GLCM texture analysis were performed to discriminate REC and RFS groups. In comparison with RFS, REC showed significant differences in perimeter (p = 0.006), integrated density (p = 0.021), contrast (p = 0.013), entropy (p = 0.002), homogeneity (p = 0.044) and inertia (p = 0.013). Prob*^REC^* was higher in the REC group than in the RFS group (0.52±0.45 versus 0.32 ± 0.42, p value = 0.038), while Prob*^RFS^* was higher in the RFS group compare to the REC group (0.82 ± 0.34 versus 0.62 ± 0.46, p=0.049). IDX*^REC^* was higher in the REC group than in the RFS group (0.47 ± 0.40 versus 0.24 ± 0.31, p=0.015) (Table 2).

**Table 2.** Comparison of MR analysis according to recurrence of endometrial cancer.

|  | No recurrence |  |  | Recurrence |  |  | *p-value* |
| --- | --- | --- | --- | --- | --- | --- | --- |
|  | N=88 |  |  | N=24 |  |  |  |
| Histogram analysis |  |  |  |  |  |  |  |
| Total area (mm<sup>2</sup>) | 549.69 | ± | 849.13 | 864.44 | ± | 557.58 | .089 |
| Perimeter | 89.87 | ± | 48.36 | 120.10 | ± | 39.95 | .006 |
| Roundness | 0.56 | ± | 0.29 | 0.66 | ± | 0.18 | .120 |
| Circularity | 0.66 | ± | 0.19 | 0.70 | ± | 0.15 | .289 |
| AR | 10.81 | ± | 78.61 | 1.66 | ± | 0.51 | .571 |
| Perfusion ratio | 0.46 | ± | 0.17 | 0.44 | ± | 0.19 | .654 |
| Integrated density | 232.08 | ± | 277.49 | 351.78 | ± | 199.85 | .021 |
| GLCM texture analysis |  |  |  |  |  |  |  |
| ASM | 0.01 | ± | 0.05 | 0.00 | ± | 0.00 | .238 |
| IDM | 0.16 | ± | 0.16 | 0.10 | ± | 0.07 | .068 |
| Contrast | 418.50 | ± | 577.51 | 768.34 | ± | 697.18 | .013 |
| Energy | 0.01 | ± | 0.05 | 0.00 | ± | 0.00 | .238 |
| Entropy | 5.88 | ± | 1.30 | 6.78 | ± | 0.89 | .002 |
| Homogeneity | 0.23 | ± | 0.16 | 0.16 | ± | 0.08 | .044 |
| Variance | 1086.32 | ± | 1226.39 | 1650.76 | ± | 1366.38 | .054 |
| Shade | -217499.07 | ± | 399322.36 | -300604.00 | ± | 362572.98 | .359 |
| Prominence | 102574664 | ± | 182479928 | 164848390 | ± | 197951255 | .148 |
| Inertia | 418.50 | ± | 577.51 | 768.34 | ± | 697.18 | .013 |
| Correlation | 0.47 | ± | 0.41 | 0.43 | ± | 0.37 | .680 |
| Deep learning |  |  |  |  |  |  |  |
| Probability of recurrence | 0.32 | ± | 0.42 | 0.52 | ± | 0.45 | .038 |
| Probability of RFS | 0.82 | ± | 0.34 | 0.62 | ± | 0.46 | .049 |
| Recurrence score | 0.24 | ± | 0.31 | 0.47 | ± | 0.40 | .015 |

Compared to the FIGO I-II, FIGO III-IV showed significant differences in the total area (p < 0.012), perimeter (p < 0.009), and integrated density (p = 0.004) in MR histogram and texture analysis. Histogram and GLCM texture analysis were performed to discriminate REC and RFS groups. In comparison with RFS, REC showed significant differences in perimeter (p = 0.006), and integrated density (p = 0.021), contrast (p = 0.013), entropy (p = 0.002), homogeneity (p = 0.044) and inertia (p = 0.013) Compared to the low-grade tumor, high-grade tumor showed significant differences in the total area (p < 0.009), perimeter (p = 0.003), perfusion ratio (p = 0.020) and integrated density (p = 0.015) in MR histogram and texture analysis (Figure 5).

**Figure 5.**
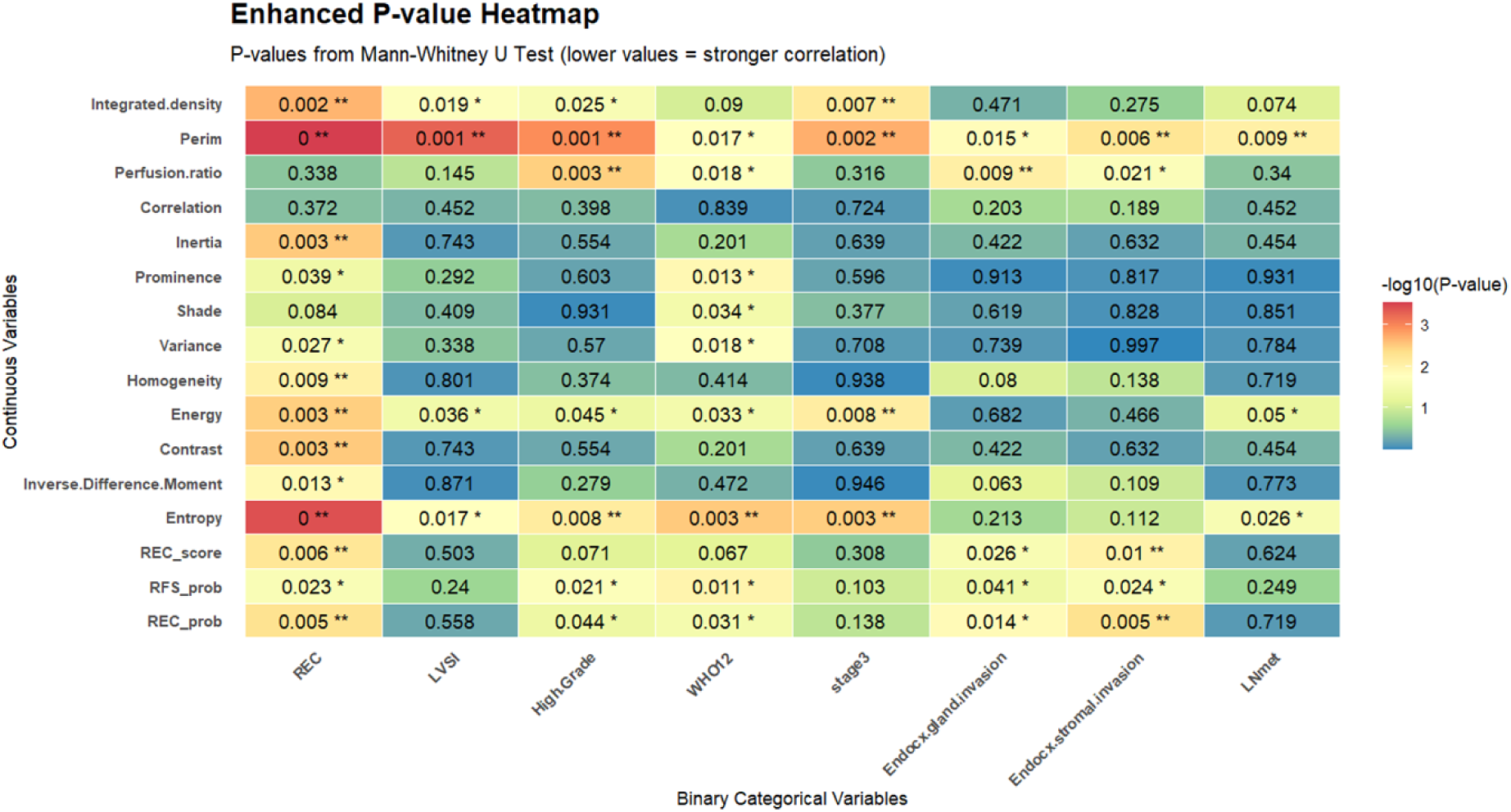
Heatmap of P-values between continuous radiomic features and binary clinical variables. The heatmap shows −log10 transformed P-values from Mann–Whitney U tests between continuous variables (e.g., Area, Entropy, REC_score) and binary clinical factors (e.g., High-grade, Stage3, LVSI). Stronger associations are shown in warmer colors. Asterisks indicate significance (* P < 0.05, ** P < 0.01, *** P < 0.001).

### IV. validation of fRCNN with imaging analysis

Receiver Operating Characteristic (ROC) Analysis of Predictive Variables. The ROC curves illustrate the diagnostic performance of three predictive variables: Integrated Density, LVI Score, and Entropy. The area under the curve (AUC) values indicate the discriminative ability of each variable, with higher AUC values representing better predictive performance. The AUC values for area, integrated density, entropy, and REC score are 0.757, 0.633, 0.705, 0.737 and 0.683, respectively. The DeLong’s test results indicate no statistically significant differences among the AUC values (p > 0.05 for all pairwise comparisons). (Figure 6).

**Figure 6.**
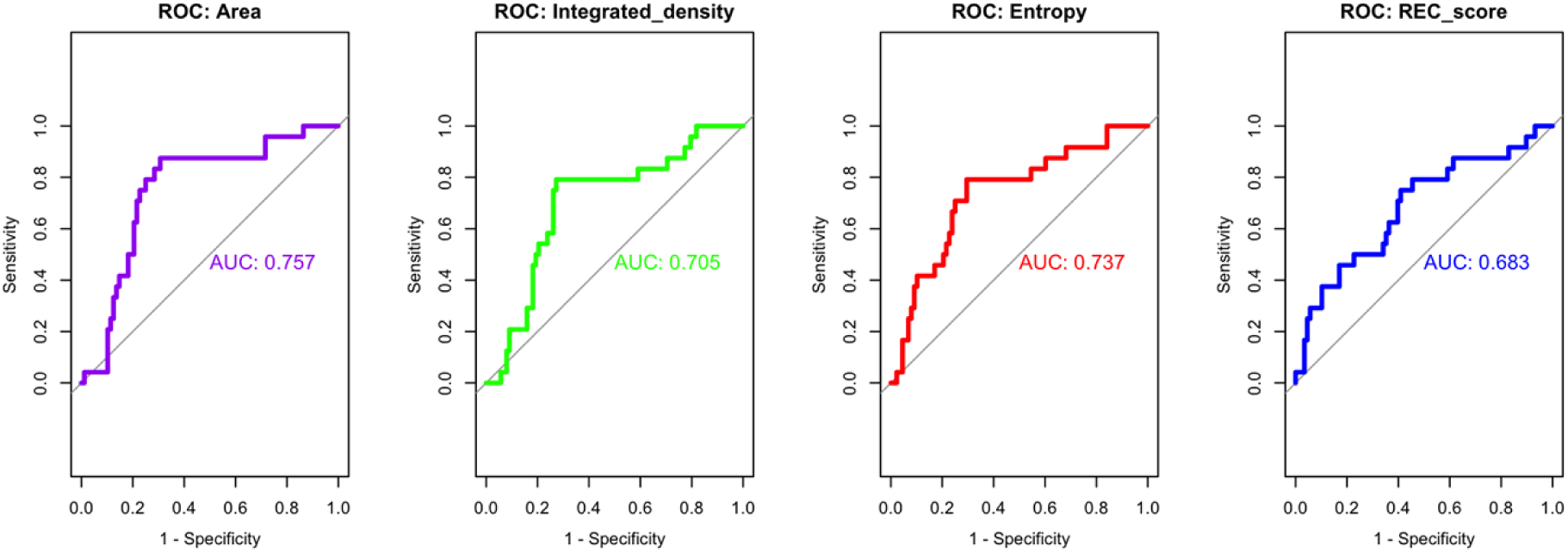
Receiver Operating Characteristic (ROC) Analysis of Predictive Variables The ROC curves illustrate the diagnostic performance of predictive variables: Integrated Density, LVI Score, and Entropy. The area under the curve (AUC) values indicate the discriminative ability of each variable, with higher AUC values representing better predictive performance.

Feature importance analysis using a Random Forest model compared REC_score from Faster R-CNN with texture-derived parameters. The Mean Decrease Gini method identified Area (4.37) as the most influential feature, while REC_score (3.36) showed similar importance to Integrated_density (2.92) and Entropy (2.81). Additionally, Perim (2.57), Homogeneity (2.34), Inertia (2.30), and inverse difference moment (2.29) were significant predictors, highlighting the role of texture uniformity in recurrence assessment (Figure 7).

**Figure 7.**
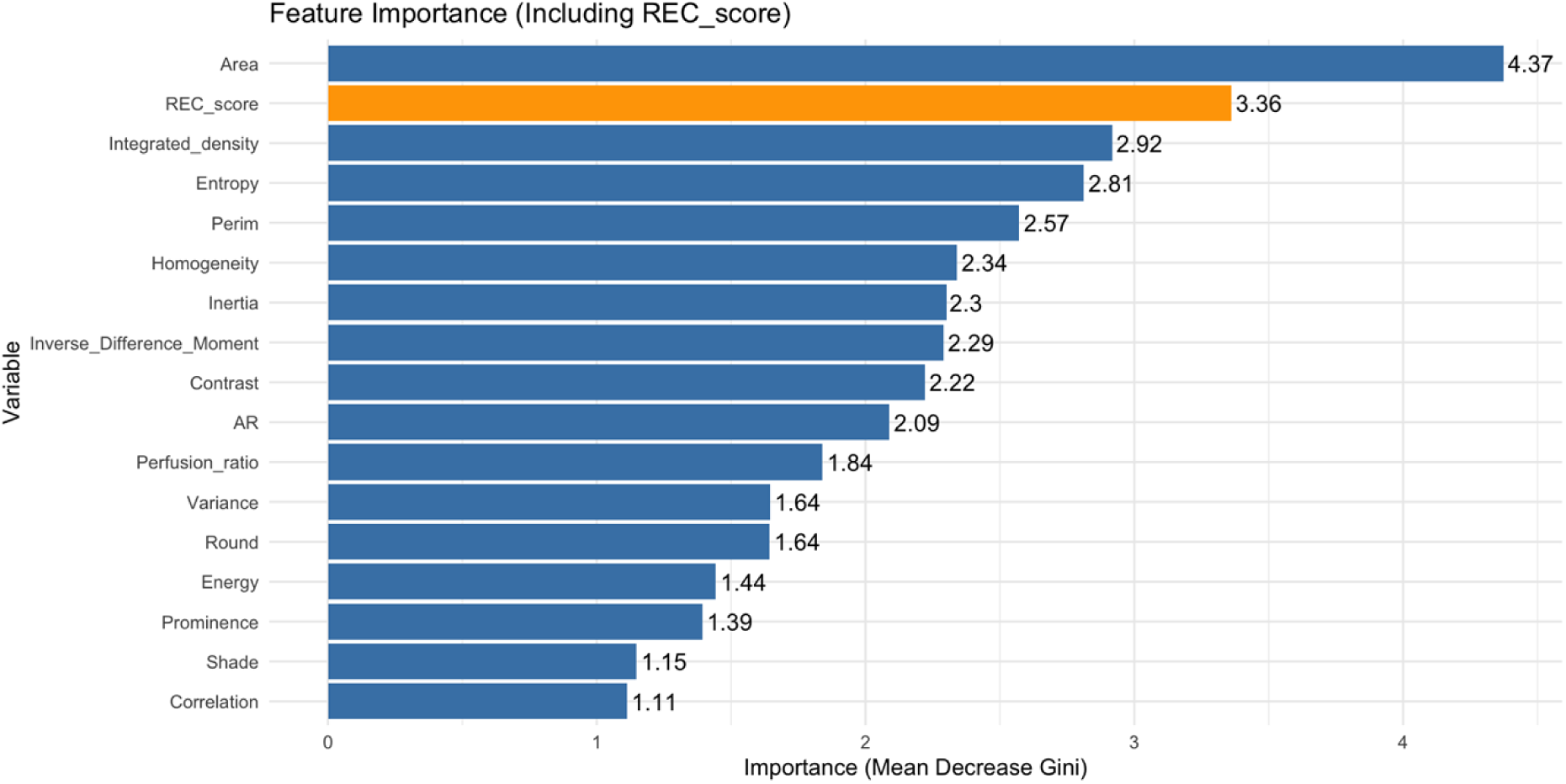
Feature importance rankings for using the Mean Decrease Gini method. The x-axis represents the importance score, indicating the contribution of each feature to the predictive model. Higher scores denote greater influence in distinguishing recurrence cases.

A correlation matrix with p-values was constructed to evaluate the relationships between REC_score and texture analysis variables. The analysis revealed strong correlations among Integrated_density, Entropy, and Perim, indicating a close association between intensity distribution and shape complexity. Additionally, Homogeneity and Inverse Difference Moment exhibited a strong correlation, confirming their role in texture uniformity. However, no statistically significant correlation was observed between REC_score and texture analysis variables. This suggests that the deep learning-based prediction score (REC_score) captures information independent of texture-based features (Figure 8).

**Figure 8.**
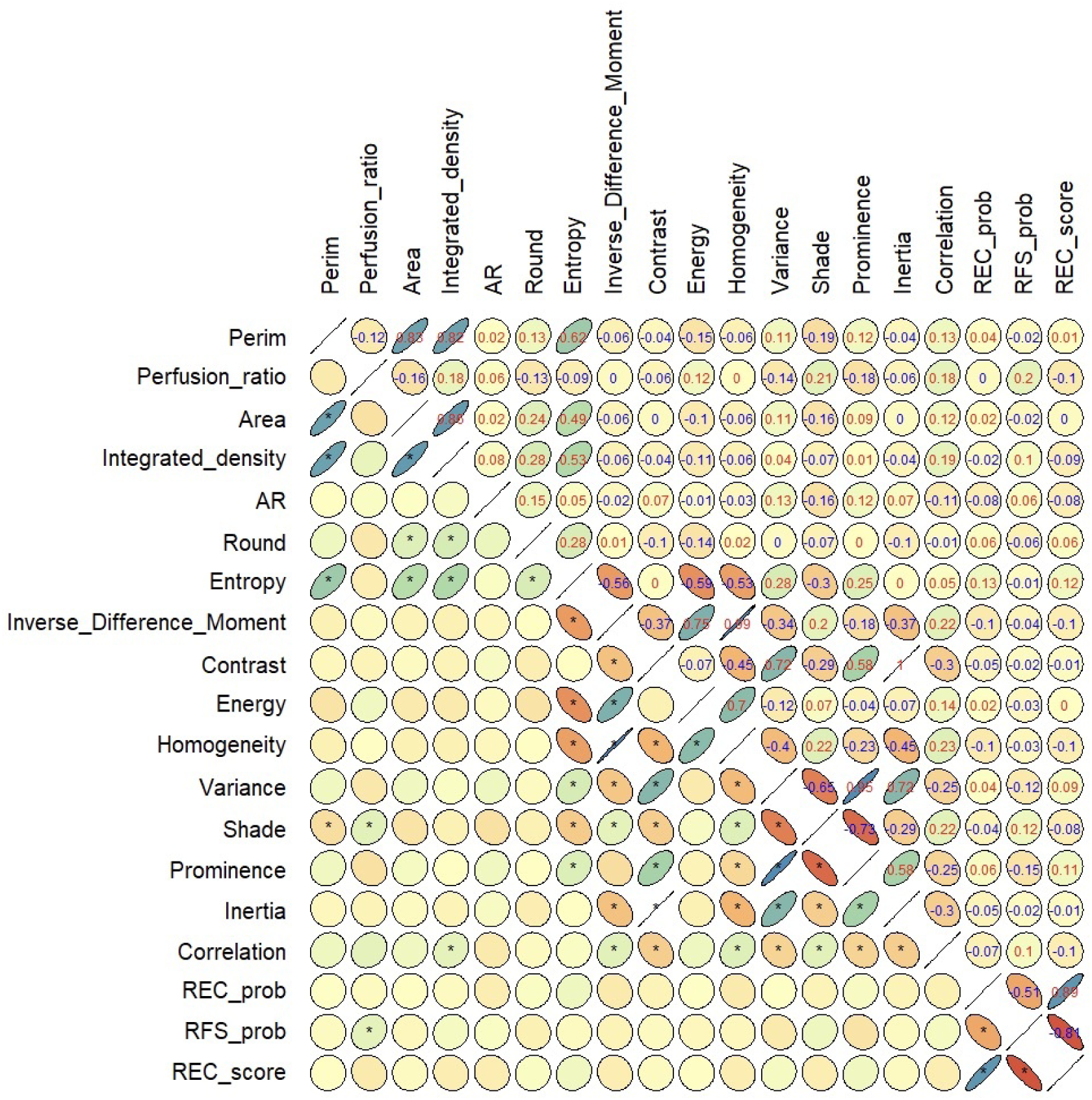
Correlation Matrix presents a correlation matrix for several features. The matrix cells display the correlation coefficients (R) between pairs of features, illustrating the strength and direction of their relationships. Asterix (*) indicate that the p-value < 0.05. Stronger correlations tend to show a more pronounced linear pattern. Markers oriented from top-right to bottom-left with blue numbers indicate positive correlations, while markers oriented from top-left to bottom-right with red numbers indicate negative correlations.

## Discussion

This study aimed to validate the Faster R-CNN model for MRI-based EC recurrence prediction by comparing its performance with traditional texture and histogram analysis. Given the growing role of artificial intelligence (AI) in medical imaging, rigorous validation of deep learning models is essential to ensure their clinical applicability and reliability. Our findings suggest that Faster R-CNN provides a novel, automated approach for EC recurrence prediction, potentially complementing or even outperforming conventional radiomic methods. However, several aspects must be considered when interpreting these results, including methodological limitations, clinical implications, and the broader impact of deep learning in oncology.

A crucial component of this study was the implementation of K-fold cross-validation, which ensured robust model validation by systematically training and testing Faster R-CNN across different partitions of the dataset [11]. This approach reduces the risk of overfitting, allowing the model to generalize better to unseen data [12, 13]. Given the limited number of EC recurrence cases, this method was essential for optimizing statistical reliability and ensuring a fair comparison with traditional texture and histogram analysis. In this study, normalized mapping of gadolinum-enhanced T1-weigted MRI were used for analysis. Normalized images are crucial for consistent image analysis, as they effectively address variations arising from different MR devices or varying image acquisition parameters within a single device [14]. By applying normalization techniques, these variations can be adjusted, resulting in a more standardized and comparable set of images for analysis.

The Faster R-CNN model demonstrated competitive predictive performance, particularly in detecting recurrence-related imaging features [9, 10]. By leveraging convolutional neural networks (CNNs), the model autonomously extracted complex spatial patterns from MRI perfusion maps, reducing human bias associated with manual feature selection. The quantitative objective index derived from Faster R-CNN provided an independent prognostic metric, reinforcing the feasibility of AI-driven recurrence prediction in EC.

Despite these advantages, potential biases within deep learning models must be acknowledged. Factors such as dataset composition, class imbalance, and variations in MRI acquisition parameters can influence model performance. Ensuring diverse, high-quality training data remains a key challenge for translating deep learning models into routine clinical practice [15].

Traditional radiomic methods, including gray-level co-occurrence matrix (GLCM)-based texture analysis and histogram analysis, have been widely used for tumor characterization and prognostication[2, 16]. These techniques capture tumor heterogeneity, intensity distribution, and spatial organization within MRI scans but are inherently limited by feature selection biases and the need for expert-driven parameter tuning. One of the most useful texture analysis parameters related to EC recurrence was entropy. Entropy refers to the amount of image information required for image compression and is a measure of the change in image intensity available to reflect intratumor heterogeneity across the entire tumor volume [17].

In contrast, Faster R-CNN offers a fully automated approach to feature extraction and classification, potentially overcoming the subjectivity associated with manual texture analysis [7]. The model’s ability to analyze high-dimensional imaging data without prior assumptions enhances its adaptability across different patient populations. Our study found that Faster R-CNN’s performance was comparable to or exceeded that of texture-based approaches, suggesting that deep learning models can complement or replace traditional radiomic techniques in recurrence risk assessment.

However, integrating deep learning into medical imaging workflows presents challenges related to model interpretability [18]. Unlike radiomic features that can be directly linked to biological phenomena, CNN-derived features are abstract and difficult to correlate with known histopathological characteristics. Future research should focus on explainable AI techniques to bridge this interpretability gap, ensuring that deep learning outputs can be meaningfully integrated into clinical decision-making.

Accurate prediction of EC recurrence is crucial for personalized treatment strategies, including enhanced surveillance, tailored adjuvant therapy, and informed patient counseling. Faster R-CNN offers an automated, quantitative approach to assessing recurrence risk, complementing existing histopathological and molecular biomarkers. Its adaptability to MRI-based evaluations aligns with the shift toward non-invasive precision oncology. Unlike biopsy-based methods, which are prone to sampling errors and procedural risks, AI-driven imaging analysis provides a comprehensive tumor assessment, capturing intratumoral heterogeneity more effectively. This advantage can improve prognostic accuracy and refine patient stratification for optimal treatment pathways. However, challenges remain before AI models like Faster R-CNN can be fully integrated into clinical practice. These include regulatory approvals, standardization of deep learning workflows, and ensuring model interpretability for clinicians. Future studies should focus on large-scale, multi-center validation to establish the generalizability of Faster R-CNN in diverse clinical settings.

Several limitations must be considered when assessing the clinical applicability of the Faster R-CNN model. First, dataset size and class imbalance may affect predictive performance. Although K-fold cross-validation was employed to mitigate bias, further validation with larger, multi-institutional datasets encompassing diverse imaging protocols and patient demographics is necessary. Second, variability in MRI acquisition parameters can impact model performance. Differences in MRI scanners, contrast agents, and imaging sequences may lead to inconsistent image quality, necessitating standardized imaging protocols to enhance reproducibility. Third, the model was trained and tested on a single dataset, which may limit its real-world applicability. External validation with independent datasets is essential to confirm its robustness and clinical relevance. Fourth, model interpretability remains a key concern. Since Faster R-CNN functions as a “black box,” it is challenging to determine which imaging features contribute to its predictions. Implementing explainable AI (XAI) techniques, such as Grad-CAM, could improve transparency and foster clinician trust. Lastly, overfitting remains a potential risk, particularly with small datasets. Strategies such as data augmentation, optimized network architectures, and larger, more diverse datasets are needed to enhance the model’s generalizability and clinical utility.

Despite several limitations, this study offers several advantages in the field of deep learning and MRI-based endometrial cancer (EC) recurrence prediction. First, this study systematically validates the Faster R-CNN model, comparing it with traditional radiomics techniques such as texture and histogram analysis. This approach enhances the reliability of deep learning-based EC recurrence prediction and establishes a robust evaluation framework. Second, this study highlights the importance of optimizing imaging acquisition techniques by assessing the relative significance of different MRI acquisition sequences. Standardizing imaging protocols can improve the accuracy of deep learning models and increase their clinical applicability. In conclusion, this study validates the feasibility of Faster R-CNN for MRI-based EC recurrence prediction, emphasizing the need for optimized imaging acquisition techniques. By analyzing the relative importance of different MRI sequences, this study contributes to improving the reliability and standardization of deep learning-based EC prediction.

## Notes

Funding disclosures: This research was supported by the Korean National Research Foundation (NRF) grants (2020R1I1A3067073) and Korean Medical Device Development Fund (RS-2023-00245686).

### Competing Interest Statement

The authors have declared no competing interest.

## References

1. Amant F, Moerman P, Neven P, Timmerman D, Van Limbergen E, Vergote I. Endometrial cancer. Lancet. 2005;366(9484):491–505. doi: 10.1016/S0140-6736(05)67063-8. PubMed PMID: 16084259.

2. Castellano G, Bonilha L, Li LM, Cendes F. Texture analysis of medical images. Clin Radiol. 2004;59(12):1061–9. Epub 2004/11/24. doi: 10.1016/j.crad.2004.07.008. PubMed PMID: 15556588.

3. Koskas M, Amant F, Mirza MR, Creutzberg CL. Cancer of the corpus uteri: 2021 update. Int J Gynaecol Obstet. 2021;155 Suppl 1(Suppl 1):45-60. doi: 10.1002/ijgo.13866. PubMed PMID: 34669196; PubMed Central PMCID: PMCPMC9297903.

4. Maheshwari E, Nougaret S, Stein EB, Rauch GM, Hwang KP, Stafford RJ, et al. Update on MRI in Evaluation and Treatment of Endometrial Cancer. Radiographics. 2022;42(7):2112–30. Epub 20220826. doi: 10.1148/rg.220070. PubMed PMID: 36018785.

5. Makker V, MacKay H, Ray-Coquard I, Levine DA, Westin SN, Aoki D, et al. Endometrial cancer. Nature Reviews Disease Primers. 2021;7(1):88. doi: 10.1038/s41572-021-00324-8.

6. Lundervold AS, Lundervold A. An overview of deep learning in medical imaging focusing on MRI. Zeitschrift für Medizinische Physik. 2019;29(2):102–27. doi: 10.1016/j.zemedi.2018.11.002 PMID - 30553609.

7. Ren S, He K, Girshick R, Sun J. Faster R-CNN: Towards Real-Time Object Detection with Region Proposal Networks. IEEE Trans Pattern Anal Mach Intell. 2017;39(6):1137–49. Epub 20160606. doi: 10.1109/TPAMI.2016.2577031. PubMed PMID: 27295650.

8. Lu Y, Yu Q, Gao Y, Zhou Y, Liu G, Dong Q, et al. Identification of Metastatic Lymph Nodes in MR Imaging with Faster Region-Based Convolutional Neural Networks. Cancer Res. 2018;78(17):5135–43. Epub 20180719. doi: 10.1158/0008-5472.CAN-18-0494. PubMed PMID: 30026330.

9. Feng M, Zhao Y, Chen J, Zhao T, Mei J, Fan Y, et al. A deep learning model for lymph node metastasis prediction based on digital histopathological images of primary endometrial cancer. Quantitative Imaging Medicine Surg. 2023;0(0):0-. doi: 10.21037/qims-22-220 PMID - 36915334.

10. Zheng R, Chen C, Zhu H, Mao W, Chen Y, Lin Y. A Deep Learning Classification Method for Early Endometrial Cancer on MRI Images. 2021 Ieee 6th Int Conf Signal Image Process Icsip. 2021;00:336–9. doi: 10.1109/icsip52628.2021.9688909.

11. Marcot BG, Hanea AM. What is an optimal value of k in k-fold cross-validation in discrete Bayesian network analysis? Computational Statistics. 2021;36(3):2009–31. doi: 10.1007/s00180-020-00999-9.

12. Bradshaw TJ, Huemann Z, Hu J, Rahmim A. A Guide to Cross-Validation for Artificial Intelligence in Medical Imaging. Radiol Artif Intell. 2023;5(4):e220232. Epub 20230524. doi: 10.1148/ryai.220232. PubMed PMID: 37529208; PubMed Central PMCID: PMCPMC10388213.

13. Hill BG, Koback FL, Schilling PL. The risk of shortcutting in deep learning algorithms for medical imaging research. Sci Rep. 2024;14(1):29224. Epub 20241125. doi: 10.1038/s41598-024-79838-6. PubMed PMID: 39587148; PubMed Central PMCID: PMCPMC11589829.

14. Kociolek M, Strzelecki M, Obuchowicz R. Does image normalization and intensity resolution impact texture classification? Comput Med Imaging Graph. 2020;81:101716. Epub 20200306. doi: 10.1016/j.compmedimag.2020.101716. PubMed PMID: 32222685.

15. Chen Z, Hu Z, Xie Y, Li D, Christodoulou AG. Repeatability-encouraging self-supervised learning reconstruction for quantitative MRI. Magn Reson Med. 2025. Epub 20250227. doi: 10.1002/mrm.30478. PubMed PMID: 40014485.

16. Haralick RM, Shanmugam K, Dinstein IH. Textural Features for Image Classification. Ieee Transactions Syst Man Cybern. 1973;SMC-3(6):610-21. doi: 10.1109/tsmc.1973.4309314.

17. Florindo J, Metze K. Using Non-Additive Entropy to Enhance Convolutional Neural Features for Texture Recognition. Entropy (Basel). 2021;23(10). Epub 20210927. doi: 10.3390/e23101259. PubMed PMID: 34681983; PubMed Central PMCID: PMCPMC8534779.

18. Do S. Explainable & Safe Artificial Intelligence in Radiology. J Korean Soc Radiol. 2024;85(5):834–47. Epub 20240927. doi: 10.3348/jksr.2024.0118. PubMed PMID: 39416324; PubMed Central PMCID: PMCPMC11473981.

